# Bigger is better: Thicker maize brace roots are advantageous for both strength and nitrogen uptake

**DOI:** 10.1101/2022.10.01.510439

**Authors:** Amanda Rasmussen, Lindsay Erndwein, Adam Stager, Jonathan Reneau, Erin E. Sparks

## Abstract

Plant root systems provide critical functions to enable plant survival. From anchoring the plant in the soil to finding and acquiring water and nutrients, these organs are essential for plant productivity. Despite a variety of root functions, research typically focuses on defining only one function. In this study, we explore a trade-off hypothesis, that the optimization of one root function (i.e. anchorage) may negatively impact another root function (i.e. nitrogen uptake). Previous work has demonstrated that larger roots are stronger, but may also have a diminished capacity for nutrient acquisition due to a reduced surface area to volume ratio. Using maize brace roots that had entered the soil, we show here that larger roots are both stronger and take up more nitrogen. Despite this general relationship, there are subtle trade-offs between mechanics and uptake that occur when assessing individual genotypes. These trade-offs represent an opportunity to optimize one root function without compromising other root functions. Together these data demonstrate that our original trade-off hypothesis was incorrect for maize brace roots, and that larger roots are both stronger and take up more nitrogen.

## Introduction

Maize is the highest produced grain crop in the world with a large variety of uses from human food to biodegradable plastics. Maize productivity relies on a healthy and efficient root system to anchor the plant and acquire nutrients and water. As a monocot, maize has a fibrous root system composed of embryonic (primary and seminal) and post-embryonic (lateral, crown and brace) root types (Blizard and Sparks, 2020). For the majority of the maize life-cycle, the post-embryonic roots are the dominant root type in terms of function (Hochholdinger, 2009). Of the post-embryonic roots, crown and brace roots both originate from stem nodes but differ by location with crown nodes from below-ground nodes and brace roots from above-ground nodes. Brace roots can either remain aerial (when positioned on higher nodes) or grow into the ground (when positioned on lower nodes). The different origins and destinations of these roots result in different contributions to root functions such as mechanical support, important for lodging resistance, and nutrient supply, required for growth and development.

A failure of root anchorage, referred to as root lodging, has detrimental impacts on maize production (Hostetler et al., 2022b; Tirado et al., 2021). We have previously shown that brace root phenotypes (from the roots that enter the soil) can predict the brace root contribution to anchorage and lodging susceptibility (Hostetler et al., 2022b). Specifically, we found that larger diameter roots and larger root systems indicate a lower incidence of lodging (Hostetler et al., 2022b). To define the biomechanics of individual brace roots, we used 3-point bend tests across multiple genotypes and nodes (Hostetler et al., 2022a). The fundamentals of biomechanics tell us that larger diameter roots will have a higher bending strength, because a larger cross-section is able to distribute more force and thus withstand greater forces. Indeed, we showed a positive relationship between brace root geometry and bending strength, although there was contribution from the underlying material properties (e.g. cell wall composition or structure) (Hostetler et al., 2022a). Collectively, larger diameter brace roots are stronger at the individual root and root system level, suggesting that larger roots are advantageous for lodging-resistance.

Another key root function, nutrient uptake, occurs through surface-expressed transporters before redistribution internally (usually called translocation) to other plant tissues and organs (Geiger, 2009). Plants can take up both organic and inorganic nitrogen forms and each have their own specialised transporter families that can be classified as high (very specialised) or low (more generalist) affinity (Dechorgnat et al., 2019; Fan et al., 2017; Rentsch et al., 2007). While organic nitrogen uptake can occur, for high-input growth systems that are typical of maize production in many parts of the world, inorganic nitrogen forms (e.g. ammonium or nitrate) are typically accepted as the dominant sources of fertiliser (Garnett et al., 2015; Vinall et al., 2012). While for some species the lower energy requirement for uptake of organic nitrogen or ammonium sources result in higher preferences for these sources (Brackin et al., 2015; Kronzucker et al., 1997; Trépanier et al., 2009; Wang and Macko, 2011), there is very little known about maize root preferences or the uptake ability of post-embryonic brace roots. A study in hydroponics demonstrated roots of 21-day old plants (presumably seminal, or potentially crown roots at this stage of development) took up almost twice the amount of ammonium compared to nitrate (Garnett et al., 2015). More recently a study fed N15 as Urea (an organic molecule) to the soil of field-grown maize over a period of a week, meaning all roots in the soil could access the labelled nitrogen, and unsurprisingly larger root systems resulted in more labelled nitrogen in the shoot (Zhang et al., 2022). While these studies have shown there are potential preference differences between inorganic forms in young plants (Garnett et al., 2015) and that their roots can take up organic forms (Zhang et al., 2022), little is known about brace root nitrogen uptake levels or preferences at the root level in older plants.

Since the first point of entry for nutrients is at the root surface, it is generally accepted that roots with a higher surface area to volume ratio (i.e. smaller diameter roots) have a higher uptake capacity per volume (De Bauw et al., 2018; Rewald et al., 2011; Santner et al., 2012). This means there is the potential for larger diameter roots (which may be stronger to resist lodging) to have less nitrogen uptake per unit root volume, compared to smaller diameter roots. In this study, we set out to test the trade-off hypothesis that larger diameter roots are stronger but take up less nitrogen per volume. Surprisingly, this hypothesis was incorrect for maize brace roots and we found that larger brace roots are both stronger and can take up more nitrogen.

## Materials & Methods

### Plant Material

Maize seeds were planted in Newark, DE USA in the summer of 2019 as part of a larger field study. At 57 days after plant (dap) (approximately vegetative leaf 10 to 11 stage, V10-V11), the following maize genotypes were used to test nutrient uptake: B73, Hickory King, Hp301, Ky21 and W64A. At 146 dap (after harvest and senescence) the brace roots of the following maize genotypes were used for mechanical testing: B73, Oh43, A632, GT112, LH252, Hickory King, Hp301, Ky21 and W64A.

### 3-Point Bend Testing

Brace root biomechanics were measured as described in (Hostetler et al., 2022a). Briefly, brace roots from the whorl closest to the soil were excised from plants, trimmed to the first 20 mm closest to the stem and retained for mechanical testing. 3-point bend testing was conducted using an Instron 5960 (Norwood, Massachusetts USA) equipped with a 100 N load cell (Instron 2530 Series static load cell, Norwood, Massachusetts USA). Samples were positioned on a custom machined 17 mm fixture and adjusted for midpoint loading with the load cell anvil in contact with the center of the sample. The tare was loaded to approximately 0.2 N and the gauge length was reset. Samples were loaded at a displacement rate of 1 mm/min. Force-displacement data were captured with Bluehill 3 software (Instron, Norwood, Massachusetts USA). Testing was halted upon sample fracture, which was defined as the first crack in the brace root section. Ultimate Load refers to the highest force the sample withstood without failure; Break Load refers to the force upon fracture. Ultimate Load and Break Load were extracted using Bluehill 3 software.

### Plant Excavation and Analysis

For brace root nutrient uptake analyses, a root ball of approximately 0.75 m in diameter was excavated with a shovel and placed in a 5-gallon bucket of water. Roots were washed with tap water to remove excess soil and immediately placed in the Plant Spinner for imaging of shoots and roots. After imaging, brace roots that had entered the soil were excised and analysed for uptake of different sources of nitrogen. Shoots were cut from the root system and hung to dry in a greenhouse for 48 to 50 days before measuring shoot dry biomass.

### Biomass Measurements & Phenotyping

The Plant Spinner acquires 360° images of maize roots and shoots simultaneously. The imaging platform was constructed of an open aluminium frame with top mounted turn-table composed of a clamp, lid to a 5-gallon bucket and stepper motor (**Figure S1**). A white sheet was hung behind the shoots, and a white poster board placed behind the roots for contrast. A Point Grey FLIR was used for root imaging, and a RealSense SDK2.0 was used for shoot imaging. Plants were rotated 360° and images captured at every degree. For roots, images from every 40° (9 images per plant) were used for analysis. For shoots, one image with the broadest view of the shoot was used for analysis. Images were first masked using the Image Segmenter App in MATLAB (R2020a). Within the Image Segmenter App masks were created with the GraphCut option and manual selection of foreground and background pixels. Masked images were exported as black and white before import into FIJI (ImageJ Version 2.1.0; Schindelin et al., 2012). In FIJI, the images were first made binary and then pixel area was calculated using the Analyze Particles function. Pixel areas were converted by the following scales: shoot images, 1in^2^ = 25px^2^; root images, 1in^2^ = 1800px^2^. Data were then converted to metric with 1in^2^ = 6.54cm^2^. The images of excised roots from the nutrient analysis were analysed using the SmartRoot FIJI (ImageJ Version 2.0.0) add-in (Lobet et al., 2011) for diameter.

### Soil Analysis

During excavation of plants for nutrient uptake analysis, soil was collected from two locations: the bottom of the hole after plants were excavated and the centre of the row adjacent to the excavated plant. Soil samples were stored at 4°C until the end of the collection day. Soil samples were dried in a 57°C oven and then submitted to the University of Delaware Soil Testing Lab for total nitrogen and total carbon analysis via combustion.

### Nitrogen Uptake

To determine the nitrogen uptake capacity of brace roots that entered the soil, each root was incubated for 30 minutes in 15 mL of solution containing ammonium sulphate, potassium nitrate and glycine with one of the following labelled in each (^15^NH_4_)SO_4_, K^15^NO_3_, or ^15^N-glycine. Total nitrogen in each treatment was 3 mM with 1 mM of the labelled compound (10% enrichment) and 1 mM each of the two other unlabelled compounds. After incubation, roots were first rinsed with 10 mM KCl and then milliQ water, before photographing and oven drying at 57°C. Dried samples were ground in liquid nitrogen and 2-10 μg were weighed into tin capsules (precise weight recorded), placed in a 96 well plate and sent to UC Davis Stable Isotope lab for analysis using an Elementar Micro Cube elemental analyzer (Elementar Analysensysteme GmbH, Hanau, Germany) interfaced to a PDZ Europa 20-20 isotope ratio mass spectrometer (Sercon Ltd., Cheshire, UK). The ^15^N uptake per gram dry weight per hour was calculated for each sample.

### Statistics

All statistics and graphing were performed using R version 4.2.0 for Mac. Specifically, Analysis of Variance (ANOVA), pairwise comparisons with Tukey HSD post hoc tests and Pearson correlation analyses were performed with default parameters in R. The following R packages were used for data generation and display: ggplot2 version 3.3.6, agricolae version 1.3.5, ggpubr version 0.4.0, reshape2 version 1.4.4, dplyr version 1.0.9, cowplot version 1.1.1, corrplot version 0.92 and ggsci version 2.9.

### Data Availability

All raw data, code used to process data and analysed data are available at: https://github.com/EESparksLab/Rasmussen_et_al_2022.

## Results

### Brace root biomechanics vary by genotype and diameter

To study trade-offs between brace root mechanics and physiology, we first analysed brace root mechanics from nine maize genotypes via 3-point bend tests. Ultimate Load, which is the maximum force the brace root can withstand, varied by genotype (ANOVA p=6.92e-12; **Figure 1A**). B73 brace roots had the lowest Ultimate Load, and KY21 and LH252 had the highest Ultimate Load (**Figure 1A**). Additional mechanical properties can be extracted from 3-point bend tests such as the Break Load (**Figure S2A**). However, the relationship between Ultimate Load and Break Load was highly correlated (**Figure S2B**), therefore we have focused on Ultimate Load as a general metric for brace root mechanics.

**Figure 1.**
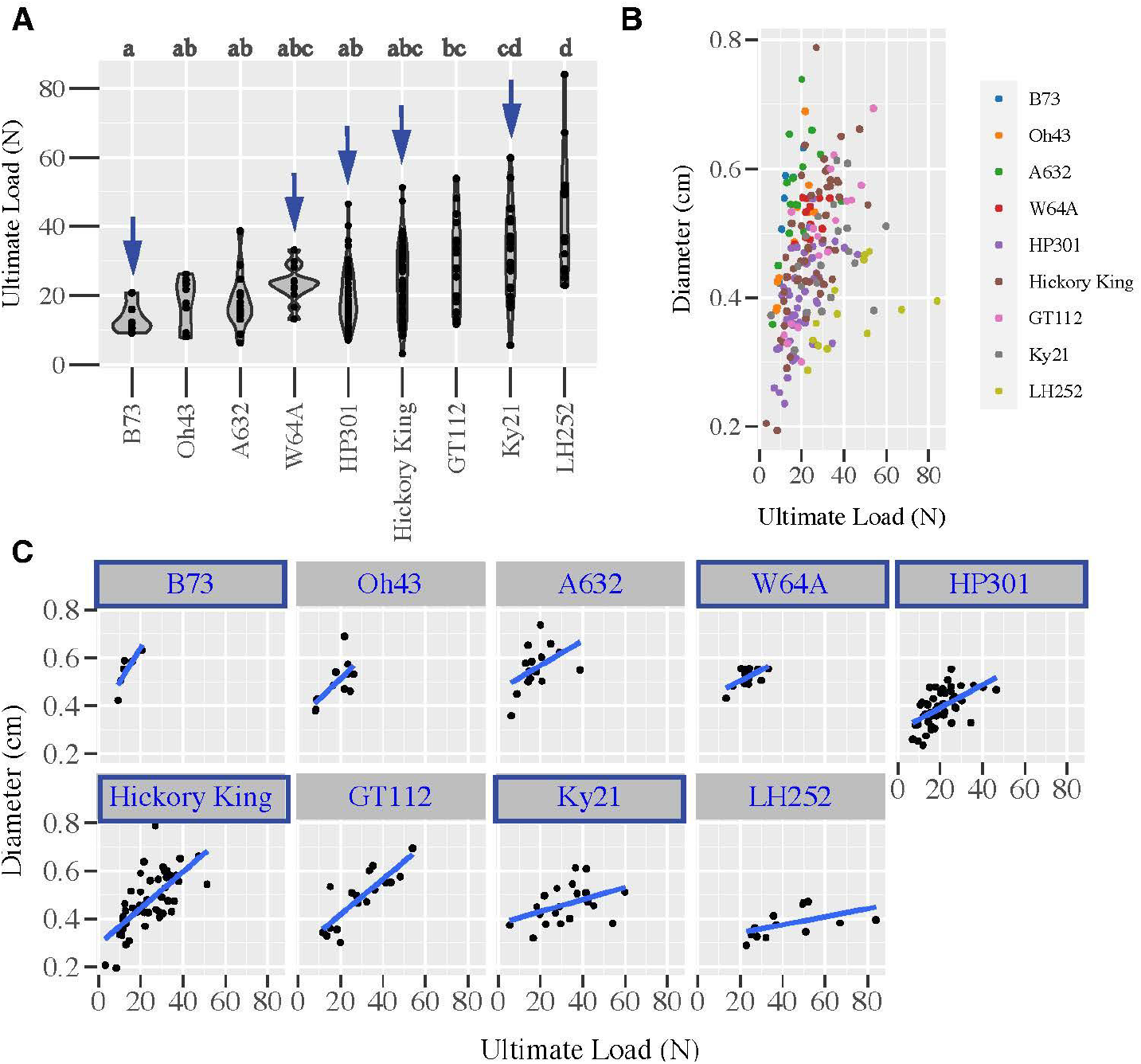
Brace root mechanics were measured by 3-point bend testing for 9 genotypes. (A) The Ultimate Load, which represents the maximum force the brace root can withstand, is variable among genotypes (p=6.92e-12). Letters indicate Tukey HSD groups. (B) There is a positive correlation between the Ultimate Load and the root diameter (r=0.32). (c) The Ultimate Load of individual genotypes are also positively correlated with brace root diameter. Genotypes marked with blue arrows and blue boxes were selected for additional analyses.

We have previously shown for three maize genotypes that, while brace root mechanics vary by whorl, genotype and reproductive stage, there is a positive correlation between brace root geometry and mechanics (Hostetler et al., 2022a). Here, brace root diameter was measured and plotted against Ultimate Load for all the genotypes combined (**Figure 1B; Table S1**) and individually (**Figure 1C; Table S1**). Consistent with previous results, there was a positive correlation between brace root mechanics and diameter (r=0.32; **Figure 1B; Table S1**), which varied by genotype (**Figure 1C; Table S1**). From these genotypes, we selected five lines with a range of brace root mechanics for detailed analysis of nutrient uptake: B73 (weakest), W64A, HP301 and Hickory king (mid-range) and Ky21 (one of the strongest).

### Shoot and root biomass allocation does not correlate with individual root mechanics

For the five selected genotypes, images of the shoots (**Figure 2A**) and roots (**Figure 2B**) were obtained immediately after excavation to estimate biomass. Shoot (**Figure 2C**) and root (**Figure 2D**) pixel area differed across the genotypes (ANOVA p=1.43e-05 and p=3.24e-05 for shoot and root respectively). There was no obvious relationship between brace root mechanics and biomass estimations. For example, two genotypes with similar, mid-range brace root mechanics (**Figure 1A)**, Hp301 and Hickory King, had the smallest and largest shoot areas respectively (**Figure 2A & C**). Dry shoot mass was also acquired (**Figure S3A**), and there was a high correlation between fresh shoot area and the dry shoot weight (**Figure S3B**) validating the pixel measure of relative biomass. The root area followed the same shoot biomass trends (**Figure 2D**), meaning genotypes with larger shoot areas also had larger root areas (r=0.70, **Figure 2C & D**; **Figure S3B**).

**Figure 2.**
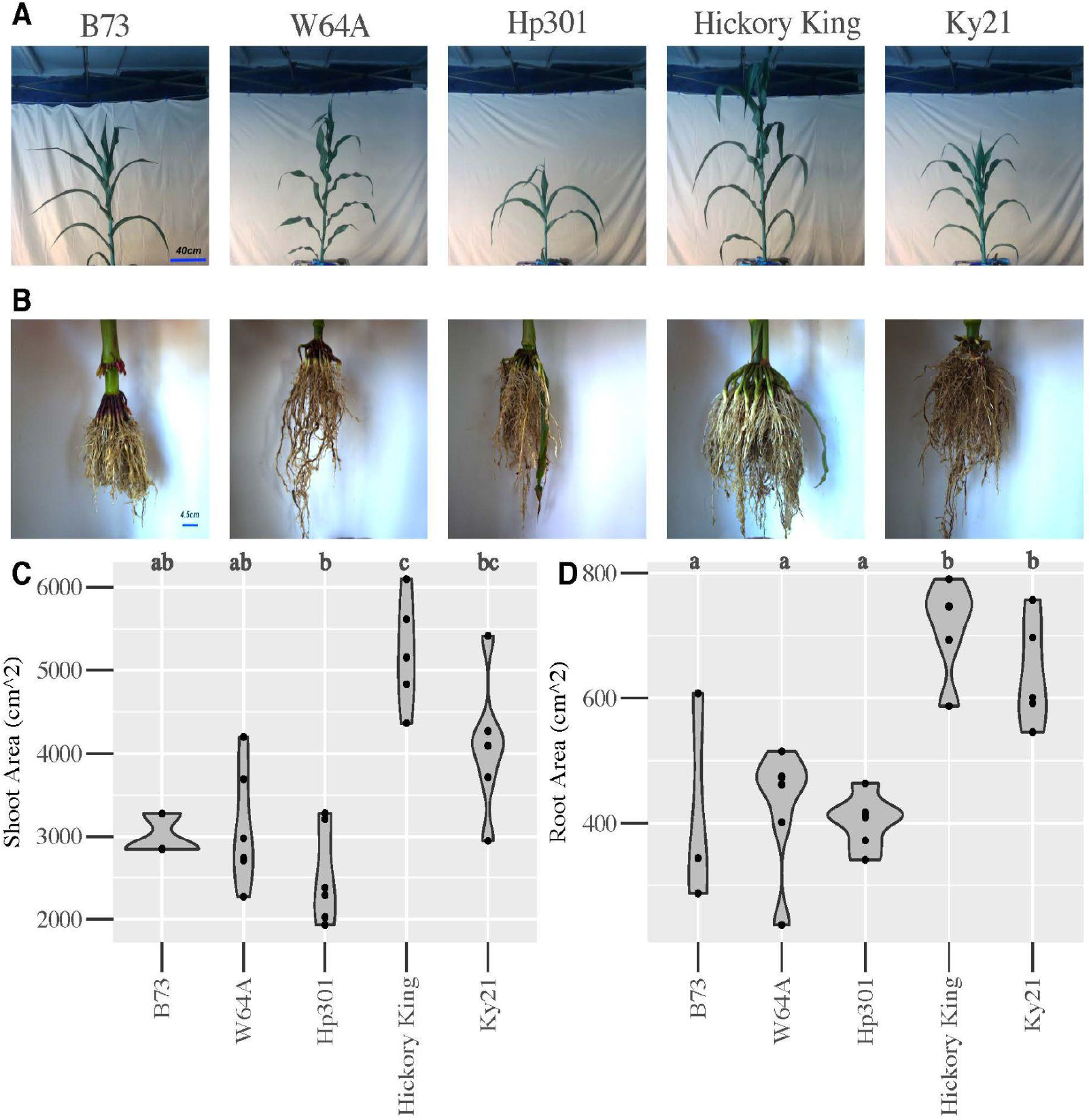
Root and shoot architecture and biomass for the selected genotypes. (A) A representative shoot image from each genotype is shown; scale bar = 40 cm. (B) The corresponding root image from each genotype is shown; scale bar = 4.5 cm. (C) Quantification of shoot area per genotype. Letters indicate Tukey HSD groups. (D) Quantification of root area per genotype. Letters indicate Tukey HSD groups. Genotypes are ordered by weakest (B73) to strongest (Ky21) brace root mechanics.

### Maize genotypes vary in the nitrogen maximum uptake capacity regardless of nitrogen source

Using the selected genotypes that vary in brace root mechanics (**Figure 1A**), we fed the ^15^N stable isotope to brace roots that entered the soil to determine nitrogen uptake capacity. Since both organic and inorganic nitrogen sources are available to plants in the field, we measured the brace root uptake of organic (glycine) and inorganic (nitrate and ammonium) sources. Considering a two-way interaction of genotype by nitrogen source. There was no statistically significant difference in the brace root uptake of glycine, ammonium, or nitrate (p=0.067; **Figure 3A**), which suggests that these plants do not have a preference for nitrogen source under these conditions. There was a significant effect of genotype on total uptake (p=0.026), with Hp301 taking up significantly more ^15^N than Hickory King (**Figure 3B**). Consistent with a lack of preference for nitrogen source, there was no interaction effect of nitrogen source by genotype (p=0.37). Thus, while there is no preference for nitrogen source, the brace roots from some genotypes uptake more nitrogen than others.

**Figure 3.**
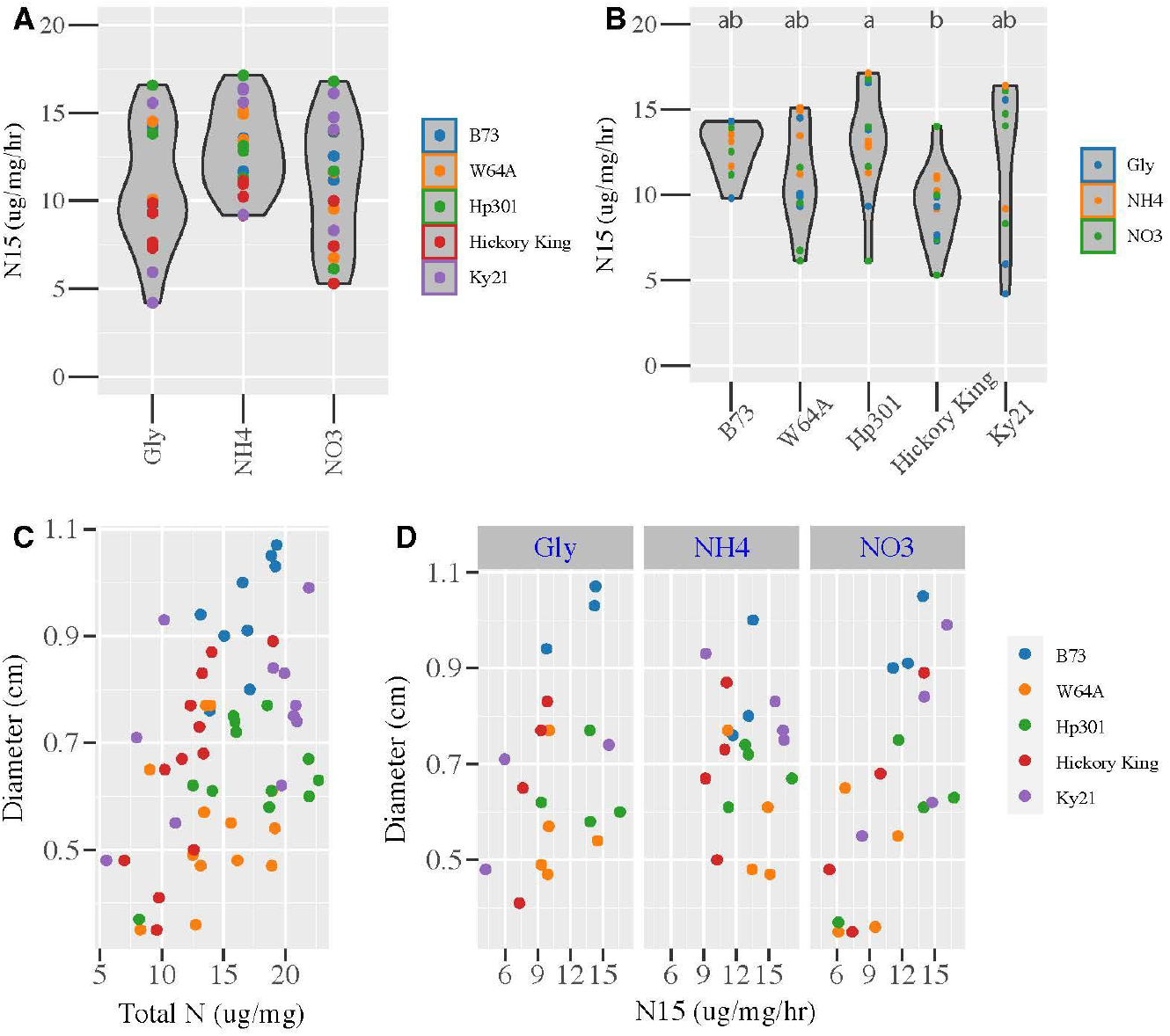
Uptake of labelled nitrogen sources. (A) There was no statistical difference in the uptake of glycine, ammonium, or nitrate among the genotypes tested. (B) Different genotypes have variable uptake of nitrogen. Letters represent Tukey HSD for genotype only. There was no interaction of genotype by nitrogen treatment (p=0.046). (C) There was a positive correlation between root diameter and total nitrogen content in each brace root (r=0.34). (D) The uptake of each nitrogen source was positively correlated with brace root diameter (r=0.35 glycine; r=-0.090 ammonium; r=0.68 nitrate). All roots received the same amount and mix of nitrogen sources with one source labelled with ^15^N. Gly =^15^N-Glycine, NH_4_ = ^15^N-labelled ammonium, NO_3_ = ^15^N-labelled nitrate.

### Nitrogen uptake increases with brace root diameter

Previous work suggests that nutrient uptake is reduced for roots with larger diameters, therefore we measured brace root diameters to characterise the relationship between nitrogen uptake and root thickness. Contrary to prior findings, there was a positive relationship between total nitrogen uptake and brace root diameter when we considered all of the genotypes together (r=0.34, **Figure 3C; Table S2**). When grouped by nitrogen source (**Figure 3D; Table S2**), differences in the relationship between uptake and diameter became apparent. Specifically, both glycine and nitrate uptake were positively correlated with brace root diameter (r=0.35 and 0.68 respectively) while ammonium was not correlated with brace root diameter (r=-0.09). Overall, these results show a positive relationship between brace root diameter and nitrogen uptake, although the nitrogen source has a variable relationship with diameter in these plants.

### Soil nutrition is uniform within and between rows

The maximum capacity of roots to take up nitrogen at any point in time is influenced by the soil conditions in which the root is growing. To determine whether heterogeneous soil conditions were influencing the ^15^N uptake results, we measured soil carbon and nitrogen levels under excavated plants and between rows (**Figure S4A**). There was no difference in nitrogen (**Figure S4B**) or carbon (**Figure S4C**) measured within and between plants for any genotype (2-way ANOVA soil acquisition point x genotype, p=0.92). There was also a high correlation between soil nitrogen and soil carbon within and between rows (**Figure S4D**; r=0.77). Lastly, there was no correlation between soil nitrogen level and uptake capacity of organic (glycine; **Figure 5A**) or inorganic (ammonium and nitrate; **Figure S5B & C**) nitrogen sources. This indicates that the soil nutrient profile is unlikely to influence differences in uptake in these plants.

### Potential trade-offs may exist at the individual genotype level

The data presented here show that larger roots have a greater mechanical strength and also take up more nitrogen. These data suggest that there was not a general trade-off between mechanics and nitrogen uptake. However, we ranked the 5 maize genotypes from weakest to strongest for average Ultimate Load, and then compared the rankings with the average total nitrogen uptake per genotype (**Table 1**). Interestingly, nuances of a potential trade-off become apparent with this analysis. Qualitatively, B73 had the weakest overall roots, but was one of the highest for nitrogen uptake. Thus, while larger roots were generally stronger and took up more nitrogen, within a given genotype there may be variation in the mechanics due to the material properties or nitrogen uptake due to transporter activity.

**Table 1.** Genotype Average Summary Statistics

|  | *Mechanics* |  | *Nitrogen Uptake* |  |  |  |
| --- | --- | --- | --- | --- | --- | --- |
|  | *Ultimate Load (N)* | *Diameter (cm)* | *Total N (ug/mg)* | *Diameter (cm)* | *UL Rank (Low-High)* | *N Rank (Low-High)* |
| *B73* | *13.46* | *0.55* | *16.68* | *0.94* | *1* | *4* |
| *HP301* | *20.22* | *0.39* | *17.10* | *0.64* | *2* | *5* |
| *W64A* | *23.44* | *0.52* | *14.26* | *0.54* | *3* | *2* |
| *Hickory King* | *24.40* | *0.48* | *12.16* | *0.65* | *4* | *1* |
| *Ky21* | *32.27* | *0.46* | *16.17* | *0.75* | *5* | *3* |

## Discussion

In this study we set out to test the trade-off hypothesis that larger diameter roots are stronger, but smaller diameter roots would take up more nitrogen per volume. While we did show that larger diameter brace roots are stronger, we were surprised to find that larger diameter brace roots also take up more nitrogen. Thus, our results refute the idea of a general trade-off hypothesis and suggest that roots, at least maize brace roots, may be optimised for both strength and nutrient supply.

The increase in nitrogen uptake scaled with root diameter such that roots approximately double in thickness were able to take up approximately twice the amount of nitrogen per root dry weight. This is consistent with our previous findings in other aerial-originating roots (Sheeran and Rasmussen, under review), but in direct contrast to embryonic (seminal) roots in wheat (Liu et al., 2022). There are a number of ways these roots could increase their net uptake including reducing nitrogen leakage back into the rhizosphere, or changing the number and type of nitrogen transporters. Given the variable relationship between individual nitrogen sources and diameter, a non-specific leakage mechanism is unlikely to explain our results. Instead, a change in the number or type of transporters is a likely explanation. Using nitrate as an example, increasing the number of nitrate transporters per surface area would increase the net maximum uptake of nitrate, or switching to low affinity transporter groups (such as NPF family) would enable the roots to take up nitrogen more generally than high-affinity transporters (such as the NRT2 family) (Dechorgnat et al., 2019; Fan et al., 2017). Further research using a combination of root anatomical imaging, transporter expression studies and Michaelis-Menten uptake kinetics is required to unravel the mechanism of increased nitrogen uptake in larger diameter maize brace roots. Additionally, a detailed analysis of nitrogen uptake in different plant species and their associated root types is required to classify root type-specific trends in nitrogen uptake.

Nitrate and ammonium (inorganic) are usually thought to be the highest available nitrogen sources in fertilised agricultural fields (Garnett et al., 2015; Vinall et al., 2012), however recent studies have highlighted that traditional methods of measuring soil nitrogen source pools may be underestimating the level of organic nitrogen sources (Brackin et al., 2015). This may mean that organic nitrogen could play larger roles in crop nutrition than previously anticipated. While an abundance of inorganic nitrogen may be predicted to drive a dominance in uptake for nitrate (also the most mobile within the soil) the energy cost of assimilation is highest for nitrate since it requires reduction to ammonium first (Crawford and Glass, 1998; Liu et al., 2013). Ammonium then needs to be bound to carbon structures, usually to Glutamine or Glutamate (Liu and von Wirén, 2017). Nitrate assimilation has been suggested to be responsible for about 25% of root carbon catabolism compared to 14% for ammonium (Taylor and Bloom, 1998), highlighting the high cost associated with nitrate despite its high abundance. Organic nitrogen sources (like glycine), on the other hand, are in an assimilation-ready state (Liu et al., 2013). Given this information we expected organic nitrogen to be favoured when roots were presented with equal availability. However, our analysis of these 5 maize genotypes showed that brace roots had no preference for nitrogen source. Part of the reason may be a result of our field having sufficient amounts of each nitrogen source, which primed the roots to express each transporter family rather than having an abundance of one type. Our field is also kept under optimum water conditions, thus maximising the availability of the nitrogen sources present. Open field light levels also mean that available energy is unlikely to be limiting assimilation of higher-cost nitrate. Under conditions where energy is limiting, such as lower light, maize may switch nitrogen preference to the lower cost sources. This may explain the doubling in ammonium uptake compared to nitrate in hydroponically grown maize in lower light levels (Garnett et al., 2015) compared to those of our open field.

While overall there is no general trade-off between support and supply, there are genotype-specific differences that could open the door to optimising both strength and nutrient uptake within a given root diameter. For example, if we rank our maize genotypes for Ultimate Load of individual roots, and compare that to the rank for nitrogen uptake, the second strongest (Hickory King) had the least nitrogen uptake while the second weakest (HP301) had the highest uptake. This outcome could be a secondary interaction as Hickory King were the biggest plants and had the most roots available to supply the shoots with nutrients, while the opposite was true for HP301. The data presented in this study were acquired at the individual brace root resolution, and thus extrapolating the strength and supply of individual roots to the whole plant remains to be determined. These nuanced patterns of strength and supply in the context of the whole plant highlight the need for future integrated, interdisciplinary studies of plant physiology.

In conclusion, our study focused on functional trade-offs under optimal conditions, and it will be very interesting to determine if resource availability (water or nutrients) will change the nitrogen uptake preferences or influence root mechanics. If under a variety of conditions, the two features remain uncoupled it will mean that root functions are advantageously linked but separable for optimisation. With increasing demand for crop productivity and a changing global climate, cross-disciplinary multi-functional studies such as these are foundational for multi-pronged plant optimisation strategies.

## Supporting information

SuppTables

SuppFigures

## Acknowledgements

We thank members of the Sparks lab, Sarah Blizard, Jeffery Sun, and Noah Ouslander for assistance in nitrogen-uptake data collection. We gratefully acknowledge members of the Sparks lab, Dr. Ashley Hostetler and Irene Ikiriko for helpful discussion and feedback on this manuscript. This work was supported by the University of Nottingham travel funds to AR and a University of Delaware Research Foundation (UDRF) award to EES. Writing of the paper occurred in person thanks to a Royal Society International Exchange grant to AR and EES.

## References

Blizard, S., Sparks, E.E., 2020. Maize Nodal Roots. Annual Plant Reviews 3, 1–24.

Brackin, R., Näsholm, T., Robinson, N., Guillou, S., Vinall, K., Lakshmanan, P., Schmidt, S., Inselsbacher, E., 2015. Nitrogen fluxes at the root-soil interface show a mismatch of nitrogen fertilizer supply and sugarcane root uptake capacity. Sci. Rep. 5, 15727.

Crawford, N.M., Glass, A.D.M., 1998. Molecular and physiological aspects of nitrate uptake in plants. Trends Plant Sci. 3, 389–395.

De Bauw, P., Vandamme, E., Lupembe, A., Mwakasege, L., Senthilkumar, K., Merckx, R., 2018. Architectural Root Responses of Rice to Reduced Water Availability Can Overcome Phosphorus Stress. Agronomy 9, 11.

Dechorgnat, J., Francis, K.L., Dhugga, K.S., Rafalski, J.A., Tyerman, S.D., Kaiser, B.N., 2019. Tissue and nitrogen-linked expression profiles of ammonium and nitrate transporters in maize. BMC Plant Biol. 19, 206.

Fan, X., Naz, M., Fan, X., Xuan, W., Miller, A.J., Xu, G., 2017. Plant nitrate transporters: from gene function to application. J. Exp. Bot. 68, 2463–2475.

Garnett, T., Plett, D., Conn, V., Conn, S., Rabie, H., Rafalski, J.A., Dhugga, K., Tester, M.A., Kaiser, B.N., 2015. Variation for N Uptake System in Maize: Genotypic Response to N Supply. Front. Plant Sci. 6, 936.

Geiger, H.H., 2009. Agronomic Traits and Maize Modifications: Nitrogen Use Efficiency. In: Bennetzen, J.L., Hake, S.C. (Eds.), Handbook of Maize: Its Biology. Springer New York, New York, NY, pp. 405–417.

Hochholdinger, F., 2009. The Maize Root System: Morphology, Anatomy, and Genetics. In: Bennetzen, J.L., Hake, S.C. (Eds.), Handbook of Maize: Its Biology. Springer New York, New York, NY, pp. 145–160.

Hostetler, A.N., Erndwein, L., Ganji, E., Reneau, J.W., Killian, M.L., Sparks, E.E., 2022a. Maize brace root mechanics vary by whorl, genotype and reproductive stage. Ann. Bot. 129, 657–668.

Hostetler, A.N., Erndwein, L., Reneau, J.W., Stager, A., Tanner, H.G., Cook, D., Sparks, E.E., 2022b. Multiple brace root phenotypes promote anchorage and limit root lodging in maize. Plant Cell Environ. 45, 1573–1583.

Kronzucker, H.J., Siddiqi, M.Y., Glass, A.D.M., 1997. Conifer root discrimination against soil nitrate and the ecology of forest succession. Nature 385, 59–61.

Liu, H., Colombi, T., Jäck, O., Westerbergh, A., Weih, M., 2022. Linking wheat nitrogen use to root traits: Shallow and thin embryonic roots enhance uptake but reduce conversion efficiency of nitrogen. Field Crops Res. 285, 108603.

Liu, X.-Y., Koba, K., Makabe, A., Li, X.-D., Yoh, M., Liu, C.-Q., 2013. Ammonium first: natural mosses prefer atmospheric ammonium but vary utilization of dissolved organic nitrogen depending on habitat and nitrogen deposition. New Phytol. 199, 407–419.

Liu, Y., von Wirén, N., 2017. Ammonium as a signal for physiological and morphological responses in plants. J. Exp. Bot. 68, 2581–2592.

Lobet, G., Pagès, L., Draye, X., 2011. A Novel Image-Analysis Toolbox Enabling Quantitative Analysis of Root System Architecture. Plant Physiol. 157, 29–39.

Rentsch, D., Schmidt, S., Tegeder, M., 2007. Transporters for uptake and allocation of organic nitrogen compounds in plants. FEBS Lett. 581, 2281–2289.

Rewald, B., Ephrath, J.E., Rachmilevitch, S., 2011. A root is a root is a root? Water uptake rates of Citrus root orders. Plant Cell Environ. 34, 33–42.

Santner, J., Smolders, E., Wenzel, W.W., Degryse, F., 2012. First observation of diffusion-limited plant root phosphorus uptake from nutrient solution. Plant Cell Environ. 35, 1558–1566.

Schindelin, J., Arganda-Carreras, I., Frise, E., Kaynig, V., Longair, M., Pietzsch, T., Preibisch, S., Rueden, C., Saalfeld, S., Schmid, B., Tinevez, J.-Y., White, D.J., Hartenstein, V., Eliceiri, K., Tomancak, P., Cardona, A., 2012. Fiji: an open-source platform for biological-image analysis. Nat. Methods 9, 676–682.

Sheeran, L. and Rasmussen, A., 2022. Aerial roots elevate indoor plant health: physiological and morphological responses of three high-humidity adapted Araceae species to indoor humidity levels. In Review.

Taylor, A.R., Bloom, A.J., 1998. Ammonium, nitrate, and proton fluxes along the maize root. Plant Cell Environ. 21, 1255–1263.

Tirado, S.B., Hirsch, C.N., Springer, N.M., 2021. Utilizing temporal measurements from UAVs to assess root lodging in maize and its impact on productivity. Field Crops Res. 262, 108014.

Trépanier, M., Lamy, M.-P., Dansereau, B., 2009. Phalaenopsis can absorb urea directly through their roots. Plant Soil 319, 95–100.

Vinall, K., Schmidt, S., Brackin, R., Lakshmanan, P., Robinson, N., 2012. Amino acids are a nitrogen source for sugarcane. Funct. Plant Biol. 39, 503–511.

Wang, L., Macko, S.A., 2011. Constrained preferences in nitrogen uptake across plant species and environments. Plant Cell Environ. 34, 525–534.

Zhang, P., Yan, Y., Gu, S., Wang, Y., Xu, C., Sheng, D., Li, Y., Wang, P., Huang, S., 2022. Lodging resistance in maize: A function of root–shoot interactions. Eur. J. Agron. 132, 126393.

