## Supplementary material for "Bigger is better: Thicker maize brace roots are advantageous for both strength and nitrogen uptake": SuppFigures

### Supplemental Information

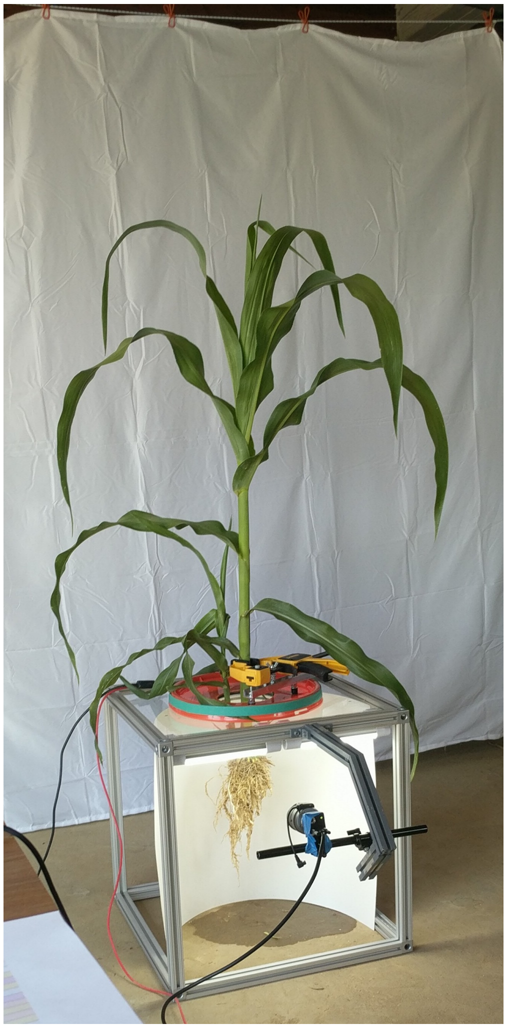

Figure S1: The Plant Spinner is composed of an aluminium framed box, a top-mounted clamp, and rotary component. Plants were placed in the Plant Spinner and images of roots and shoots were acquired at 1-degree intervals.

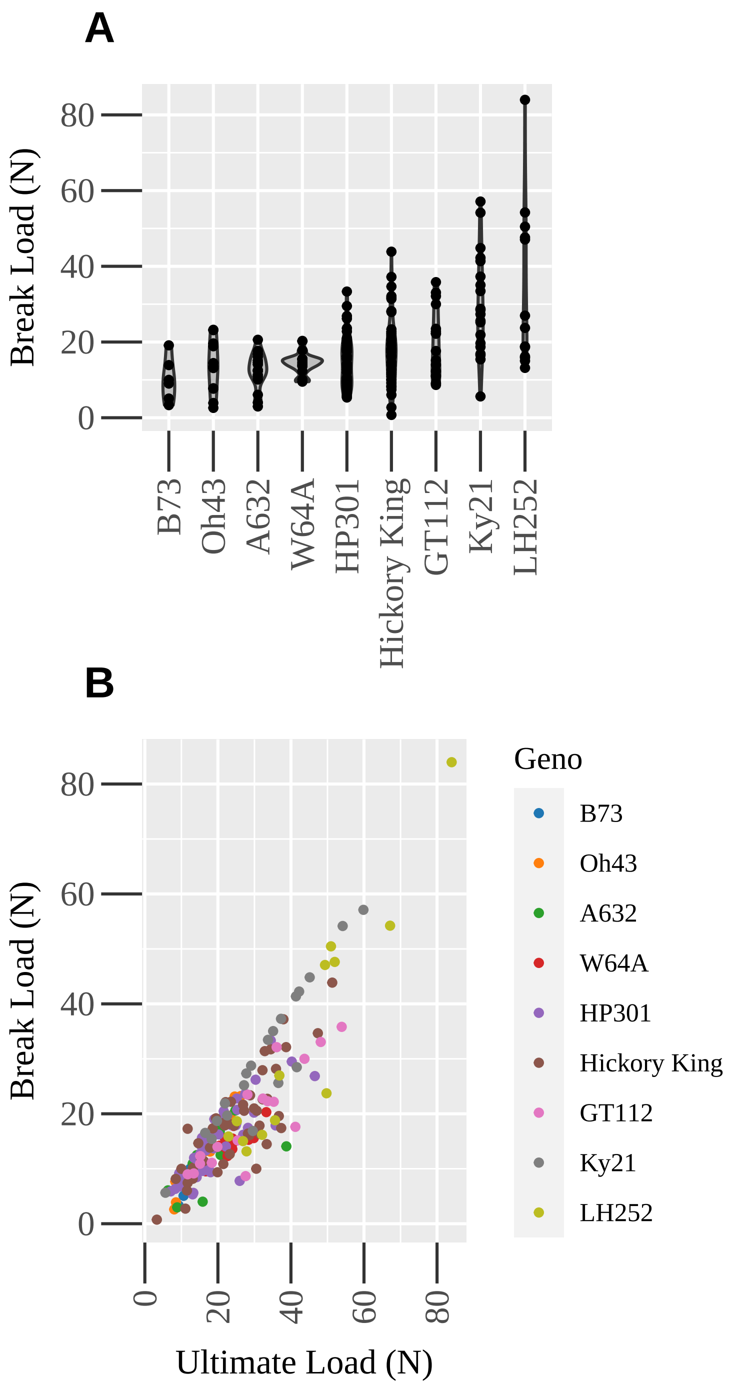


Figure S2: Brace root biomechanics were measured by 3-point bend testing for 9 inbred genotypes. The (A) Break Load was variable by genotype and (B) was highly correlated with the Ultimate Load.

**A**

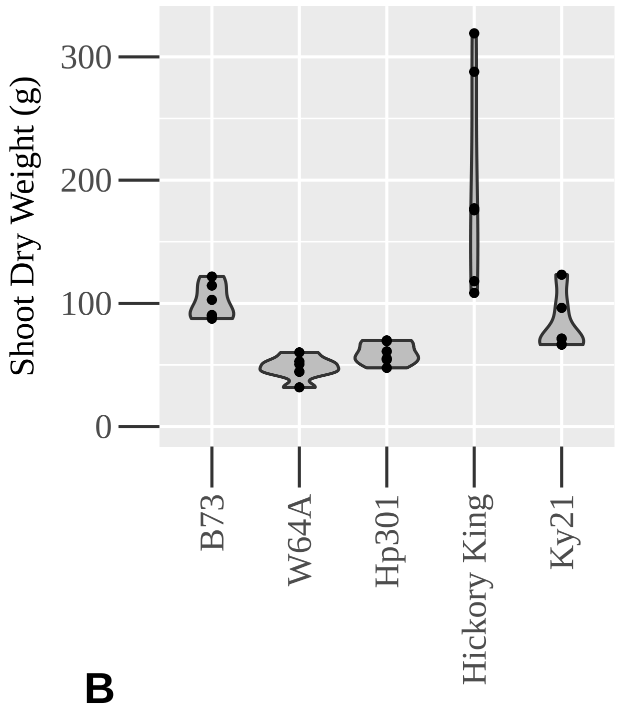


**B**

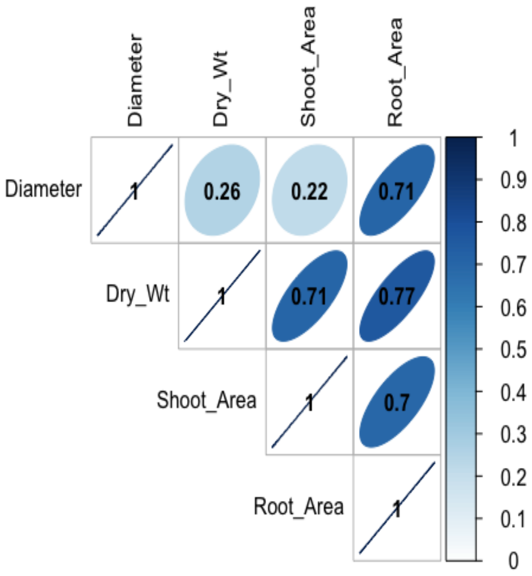


Figure S3: (A) Shoot dry weights were collected from the plants analyzed for nitrogen uptake. Hickory King dry weight was significantly higher than the other genotypes by ANOVA and Tukey HSD p-value <0.01. (B) There was high correlation among the plant biomass characteristics.

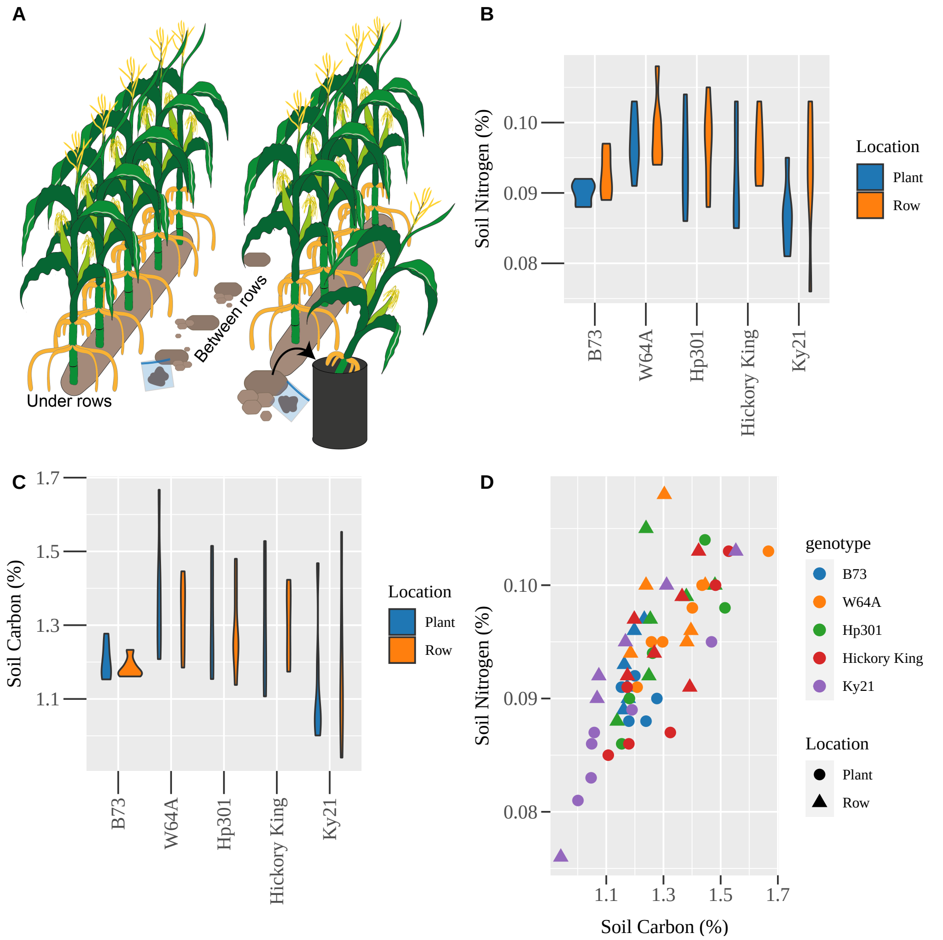


Figure S4: Soil carbon and nitrogen levels. (A) Illustration of sample collection. (B) There was no difference in soil nitrogen (ANOVA p=0.92) or (C) soil carbon (ANOVA p=0.93) between collection from below plants (blue) and between rows (orange) within a genotype. (D) A high positive correlation (r=0.76) between percent nitrogen and percent carbon suggests that nutrients are present as organic matter rather than residual inorganic fertiliser.

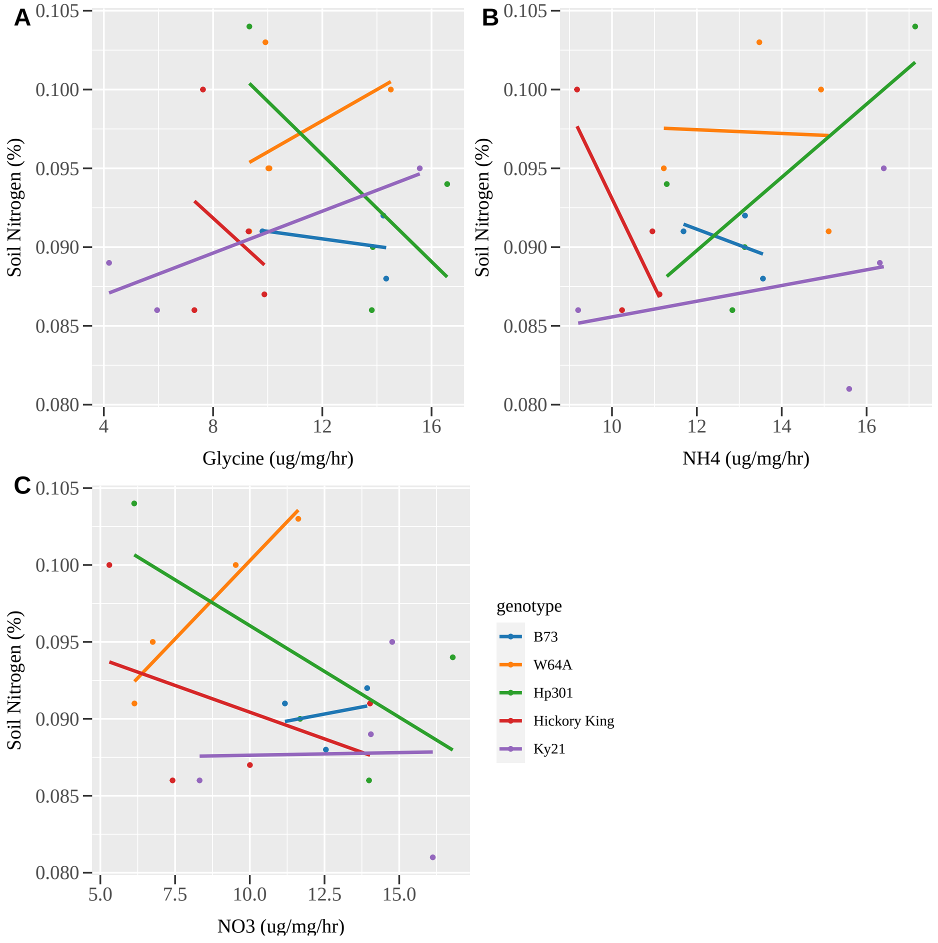


Figure S5: Soil nitrogen by brace root nitrogen uptake. There is no correlation between soil nitrogen levels and root nitrogen uptake of (A) glycine, (B) ammonium, or (c) nitrate across the five lines measured.
